# Translating DNA origami Nanotechnology to Middle School, High School, and Undergraduate Laboratories

**DOI:** 10.1101/2022.09.15.508130

**Authors:** P. E. Beshay, A. Kucinic, N. Wile, P. Halley, L. Des Rosiers, A. Chowdhury, J. L. Hall, C. E. Castro, M. W. Hudoba

## Abstract

DNA origami is a rapidly emerging nanotechnology that enables researchers to create nanostructures with unprecedented geometric precision that have tremendous potential to advance a variety of fields including molecular sensing, robotics, and nanomedicine. Hence, many students could benefit from exposure to basic knowledge of DNA origami nanotechnology. However, due to the complexity of design, cost of materials, and cost of equipment, experiments with DNA origami have been limited mainly to research institutions in graduate level laboratories with significant prior expertise and well-equipped laboratories. This work focuses on overcoming critical barriers to translating DNA origami methods to educational laboratory settings. In particular, we present a streamlined protocol for fabrication and analysis of DNA origami nanostructures that can be carried out within a 2-hour laboratory course using low-cost equipment, much of which is readily available in educational laboratories and science classrooms. We focus this educational experiment module on a DNA origami nanorod structure that was previously developed for drug delivery applications. In addition to fabricating nanostructures, we demonstrate a protocol for students to analyze structures via gel electrophoresis using classroom-ready gel equipment. These results establish a basis to expose students to DNA origami nanotechnology and can enable or reinforce valuable learning milestones in fields such as biomaterials, biological engineering, and nanomedicine. Furthermore, introducing students to DNA nanotechnology and related fields can also have the potential to increase interest and future involvement by young students.

**GRAPHICAL ABSTRACT:** 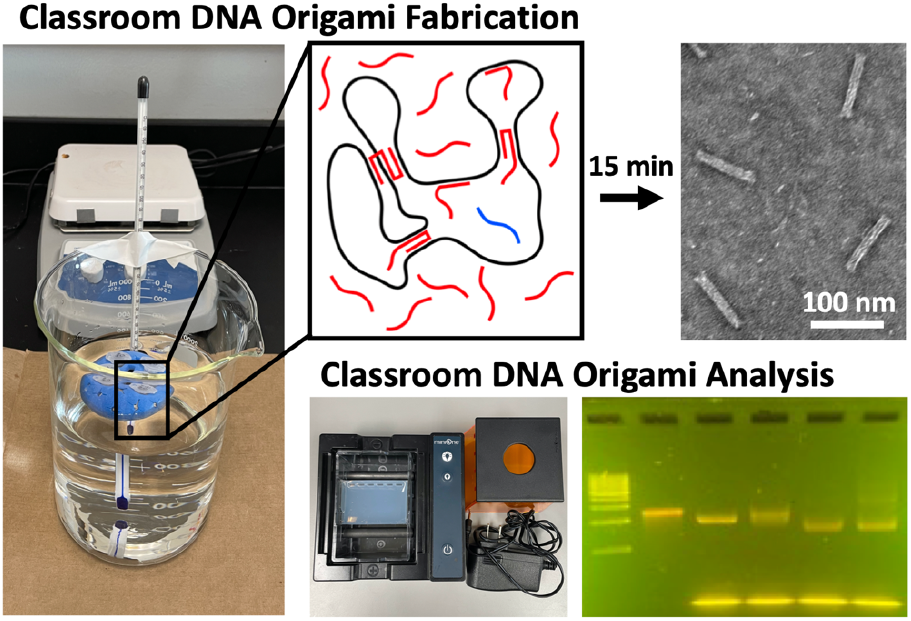

## INTRODUCTION

DNA origami is a rapidly emerging technology that has demonstrated tremendous promise for applications including molecular sensing, nanorobotics, and nanomedicine^1–5^. In this approach, a long single-stranded DNA (ssDNA) ‘scaffold’ strand is folded into a compact structure via DNA base-pairing interactions with many shorter ssDNA ‘staple’ strands, allowing researchers to create nanostructures with unprecedented geometric precision via molecular self-assembly^6–8^. Due to the complexity of design and the cost of materials and equipment, DNA origami studies have mainly been limited to research institutions in graduate level laboratories with significant prior expertise and well-equipped laboratories. However, many students can benefit from exposure to and basic knowledge of DNA origami nanotechnology since it is likely to impact a wide range of fields and industries. Furthermore, DNA origami can serve as an introduction to more general biomolecular engineering concepts; and given the wide range of functions that have been implemented in DNA design (e.g., mechanical deformation^9^, polymerization^10^, actuation^11^ etc.), DNA nanotechnology can be a unique way to introduce or reinforce other science and engineering concepts. Over the long-term, introducing a broad range of students to DNA origami would also have the potential to advance the field due to increased interest and involvement by young students, who may then pursue education, research, or career paths related to DNA nanotechnology^12^.

We believe that by circumventing the complexity of the design process and removing the hefty cost and infrastructure associated with DNA origami fabrication, valuable educational milestones can be achieved by young students in fields such as engineering, chemistry, physics, biology, materials science, medicine, and computer science. For example, specific learning opportunities that lie in DNA nanostructure fabrication include topics such as charge screening, mechanical deformations, conformational dynamics and free energy landscapes, nanoscale stimulus response, polymerization, and algorithmic design and assembly^13–18^. However, current DNA origami methods are not suitable for translation to classrooms, even for well-equipped instructional laboratories. DNA origami development often require days, or up to several weeks of design and optimization^19–21^. Furthermore, fabrication typically takes many hours or days and is carried out on costly PCR thermocyclers^19,22^, while the analysis of structure folding and behavior can take several hours with the most common first step analysis carried out using laboratory gel electrophoresis equipment^19,23^. These demands leave these topics out of reach for most undergraduate and high school educational laboratories and middle school science rooms.

To facilitate educational translation of DNA origami methods, here we developed a streamlined approach to introduce and carry out DNA origami fabrication in educational laboratories or classrooms with equipment that is either readily available or relatively inexpensive. The entire streamlined fabrication and analysis process can be carried out within a 2-hour lab session, or in 1 hour with additional teacher preparation, making it viable to carry out in standard laboratory class periods. We present a specific laboratory module, based on a previously published DNA origami nanostructure^3,24^, that introduces the concept and importance of charge screening during the folding process of a DNA origami nanodevice. We anticipate this work can open a door to introducing DNA origami to undergraduate, secondary, and primary school students and serve as a foundational example to stimulate additional educational translation related to DNA origami nanotechnology.

### DNA origami: Design, fabrication, and analysis overview

This year marks the 40^th^ anniversary of the original conception of making synthetic nanostructures out of DNA with the idea of building 2D or 3D lattices out of many similar copies of nucleic acid junctions^25^. In 2006, the development of scaffolded DNA origami^6^, which we refer to here as just DNA origami, took a major step in enabling more complex nanostructure geometries with a robust and versatile design and fabrication process. DNA origami is based on folding a long scaffold strand, typically ~7000-8000 nucleotides long, into a compact nanostructure through Watson-Crick^1^ base-pairing interactions^26–28^ with many, often ~150-200, shorter staple strands that are ~30-50 nucleotides long. The staple strands are designed to be piecewise complementary to the scaffold so binding pinches, or folds (hence the term origami), the scaffold into the desired shape (Fig. 1A-B). Those shapes typically consist of several dsDNA helices connected in parallel into bundles with a prescribed geometry where helices are connected to their neighbors at regular intervals by junctions (similar to Holliday junctions^29,30^) where scaffold or staples cross from one helix to the neighboring one (Fig. 1C-D). This approach allows for the fabrication of precise nanostructures with dimensions on the 5-100’s nm scale and unprecedented geometric complexity. Examples include 100 nm wide smiley faces^6^, intricate ~5-100 nm wireframe structures^31,32^, dynamic components like hinges^13,33^, sliders^33,34^, or rotors^35^, or even ~150 nm sized airplanes^21^. Here we focused our development of classroom methods for DNA origami fabrication on a previously designed nanorod structure that is in development as a drug delivery device^3^. The device is referred to as the “Horse,” as in the original publication, inspired by the concept of the Trojan Horse.

**Figure 1:**
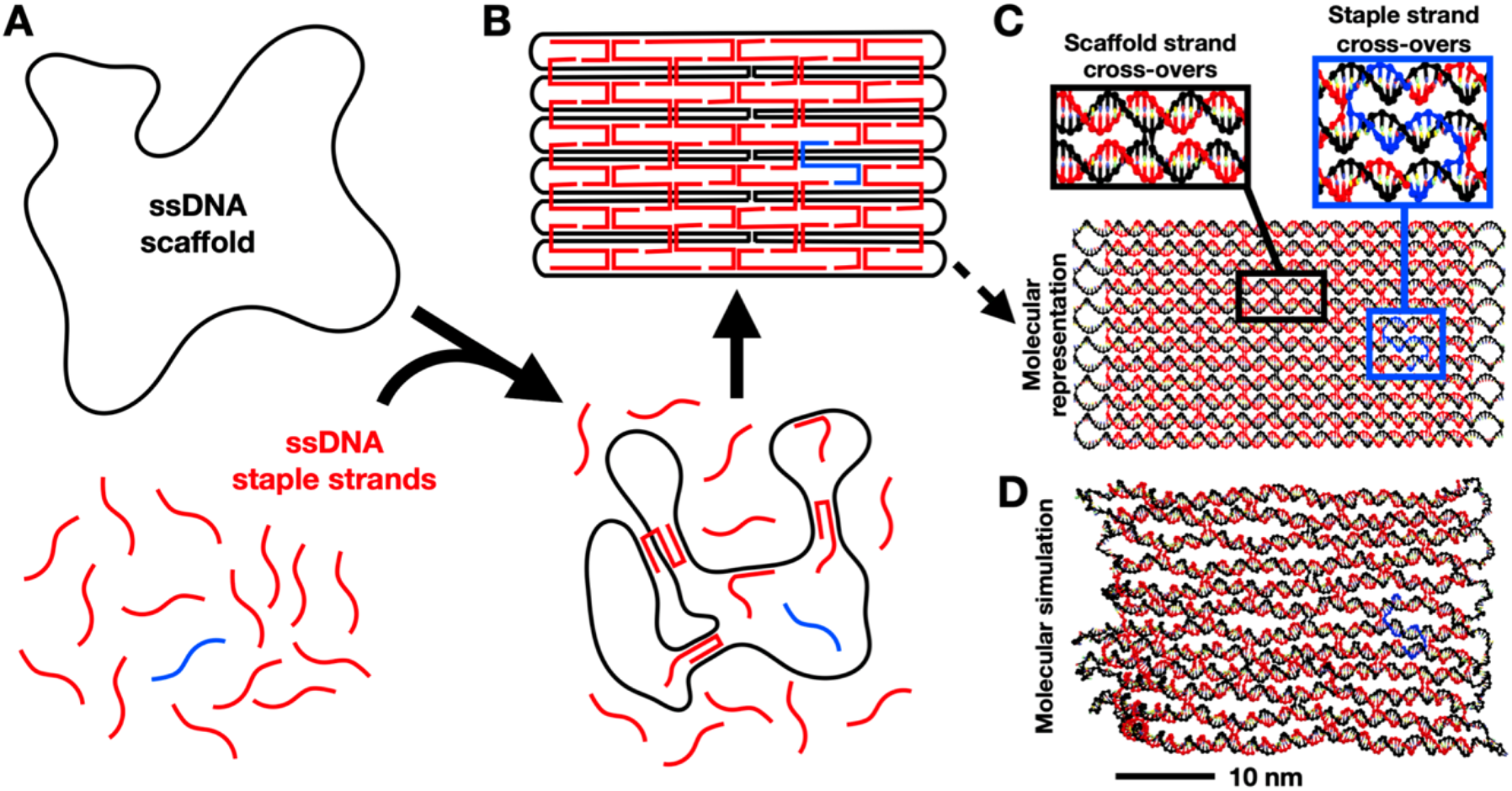
DNA origami self-assembly. A) A long (usually thousands of nucleotides) ssDNA scaffold is folded into a compact structure via base-pairing interactions with many ssDNA staple strands. The staple strands are designed to be piecewise complementary to the scaffold so they ‘pinch’ or fold the scaffold into the target shape. B) A schematic scaffold and staple strand routing design illustrates how the staples collectively hold the scaffold in the target shape, in this case an example of a rectangular plate design. Each staple, for example the blue strand, is incorporated at a specific location based on its sequence. C) A molecular model shows how the DNA helical geometry enables cross-over connections between neighboring helices, and D) a coarse-grained molecular dynamics simulation with the oxDNA model illustrates a more realistic depiction of the DNA origami rectangular plate design.

The basic DNA origami nanostructure design process^20^ follows several common steps: 1) defining the geometry (i.e., cross-section in terms of dsDNA helices and the lengths of those helices); 2) scaffold routing; 3) staple routing; and 4) staple sequence determination. These steps are typically carried out using custom computer aided design (CAD) software. The most widely used software since the development of DNA origami is caDNAno^36^, and a number of more recent tools have led to faster, partially or fully automated, and more advanced design capabilities^21,32,37–39^. In addition, significant research over the last decade has led to a number of simulation tools^19,40–43^ that are useful to predict the structure of DNA origami. Fig. 1D illustrates a molecular simulation for the small DNA origami rectangular plate example simulated using the oxDNA coarse-grained molecular dynamics model^40,41^. The DNA origami structure of interest for this work, the Horse nanostructure, was designed in the software caDNAno^36^. Although we circumvent the design process in our educational translation by using a previously published device, this work can still serve as a basis for introducing and learning the DNA origami design process. To facilitate that we provided a schematic of the Horse structure caDNAno design in Figure S1.

Once the staples are designed, they are typically ordered from one of several commercial vendors who synthesize custom DNA oligonucleotides, and the scaffold can also be purchased from a commercial vendor or produced in a laboratory as previously described^19^. Once the scaffold and staples are obtained, the fabrication, or folding, of DNA origami structures is carried out via a temperature-controlled molecular self-assembly process. Commonly used fabrication protocols are described in detail here^19,23,44^. To fold the structure, the combination of staples strands is mixed in 10-fold excess relative to the scaffold strand, and the mixture is subjected to a thermal folding ramp that consists of three phases: a melting phase, an annealing phase, and a cooling phase. Details can vary from structure to structure and are often subject to optimization for individual structures. Generally, the thermal annealing can take many hours or up to several days. For the case of the Horse nanostructure the original fabrication thermal ramp consisted of melting at 65°C for 10 minutes, followed by slow cooling from 60°C to 25°C over the course of 17 hours, and finally rapid cooling to 4°C^3^.

After fabrication, a common first step assay to evaluate folding is agarose gel electrophoresis^19^. The well-folded compact structures typically migrate through the gel faster than misfolded structures. In most (but not all) cases the compact folded structure also runs faster than the component scaffold strand, which can be included as a reference. A sharp band that migrates on the gel faster than the scaffold is typically indicative of a well-folded structure. Gel electrophoresis can also serve as a convenient purification approach to separate well-folded from mis-folded structures and from excess staple strands. To confirm folding and quantify shape and distributions of conformations, folded structures are subjected to imaging via transmission electron microscopy (TEM) or atomic force microscopy (AFM). Protocols for TEM imaging are provided in the methods and both methods are described in detail here^19,21,45,46^.

### Key barriers to educational translation

This work focuses on eliminating major barriers that make DNA origami fabrication and experiments challenging to perform in educational laboratories and classroom environments. The barriers are primarily related to resources and time required for DNA origami. The first major barrier of resources is due to the equipment needs that range from possibly available (e.g., gel electrophoresis), to unlikely available (e.g., thermocyclers), or impractical (e.g., AFM or TEM) for ready access in educational settings. The barrier of time comes from all stages of the process including: 1) Design - designing DNA origami structures can take days to weeks, especially for new designers, and even more recent automated or partially automated tools take time to learn; 2) Fabrication – self-assembly reactions can take several hours to prepare and up to days to run the thermal ramps; 3) Analysis – gel electrophoresis typically requires 2-3 hours to setup and run and AFM or TEM imaging is likely impractical for most educational settings. In addition to these barriers, the shear complexity of designs and the fabrication process are a challenge to educational translation. Here we overcome these barriers to enable the hands-on introduction and use of DNA origami technology in instructional labs and classrooms.

## METHODS

### Folding DNA origami

The horse nanostructures were folded in a single-pot reaction with 200 nM single-stranded DNA oligos (Integrated DNA Technologies), 20 nM M13mp18-derived scaffolds (prepared in-house as described in Castro, et al^19^), 20 mM MgCl_2_ (unless otherwise noted), and a buffer containing 5 mM Tris, 5mM NaCl (pH 8), and 1 mM EDTA. 1.5 mL Eppendorf tubes were used for classroom folding experiments and 200 μL Eppendorf tubes were used for laboratory folding experiments. The respective folding equipment and conditions are described below for the laboratory and classroom protocols.

#### Laboratory folding

The single-pot reaction was placed into a thermocycler (Bio-Rad) first at 65°C for 5 minutes to melt the mixture and next to anneal for certain time and temperature points as described in the results and discussions in the main text. The mixture was then cooled to 4°C and placed into a refrigerator until purification and further analysis.

#### Classroom folding

The single-pot reaction was placed in a water bath at 65°C for 5 minutes for a melting phase, then exposed to a water bath at 52.5±0.5°C for different time points as described in the results and discussion in the main text. The mixture was then cooled in an ice bath until purification.

### Purification of DNA origami

DNA origami Horse nanostructures were purified via agarose gel electrophoresis using the respective gel electrophoresis kit for the laboratory or classroom as described below.

#### Laboratory purification

Folded DNA origami nanostructures are purified via Thermo Scientific™ Owl™ EasyCast™ B1 mini gel electrophoresis kit. For the EasyCast gel, 140 mL of running buffer was created by mixing 7 mL of 10x TBE (Tris/Borate/EDTA buffer containing 45 mM boric acid, 45 mM Tris(hydroxymethyl)aminomethane base, and 1 mM (Ethylenedinitrilo)tetraacetic acid), 611 μL of 1.375 M MgCl_2_, and 132.4 mL of double-distilled water. The gel was cast by microwaving 0.62 g agarose with 61.8 g distilled water. Once the agarose is dissolved and evaporated water is replaced and 250 μL of 1.375 M MgCl_2_, and 3 μL of SYBR safe or 0.5 μg ml^-1^ ethidium bromide DNA stain was mixed. The gel was then poured and allowed to solidify. 15 μL of folded Horse structure and 3 μL of blue loading dye were pipetted into the wells of the solidified gel. Running buffer was poured into the gel rig and the gel was run at 90 V for 90 minutes in an ice water bath. The gel was then imaged on an ultra-violet light table.

#### Classroom purification

Folded DNA origami nanostructures are purified via the MiniOne agarose gel electrophoresis kit. For the MiniOne gel, 140 mL of running buffer was created by mixing 7 mL 10x TBE, 300 μL of 1.375 M MgCl_2_, and 132.7 mL of distilled water. Next, the gel was cast by microwaving 0.5 g agarose with 49.6 g distilled water. After dissolving the agarose and replacing any evaporated water, 109 μL of 1.375 M MgCl_2_, and 4 μL of GelGreen DNA stain was mixed. The gel was then poured and allowed to solidify. 8 μL of folded Horse structure and 2 μL of orange loading dye were pipetted into the wells of the solidified gel. Running buffer was poured into the MiniOne gel rig and the gel was run for 30-40 minutes at 42 V. The gel is imaged via the blue light equipped in the MiniOne gel rig with a cell phone.

### Imaging DNA origami

Purified DNA origami Horse nanostructures were suspended in the respective running buffer conditions post-gel electrophoresis at concentrations between 1 nM and 5 nM. A 4 μL sample droplet was deposited onto a plasma-treated Formvar-coated 400 mesh copper grid (Ted Pella) and incubated for 4 minutes. The droplet was wicked away on filter paper, afterwards the grid picked up a 10 μL droplet of staining solution containing 2% uranyl formate and 25 mM NaOH and then immediately wicked away. This was followed by picking up a 20 μL droplet of the same staining solution and incubating for 40 seconds before wicking away on the filter paper. The prepared samples were then dried for at least 20 minutes before imaging. The structures were imaged at the Ohio State University Campus Microscopy and Imaging Facility on a FEI Tecnai G2 Spirit TEM at 80 kV acceleration.

## RESULTS AND DISCUSSION

### DNA origami design

Designing structures using caDNAno^36^ can take days to weeks, especially for new designers. Recent automated or semi-automated tools allow for design in minutes^21,32,37–39^. These approaches still require introducing software, which can take several hours to days to present and learn to operate. We circumvent the design process by relying on the previously published Horse nanostructure^3^ (Fig. 2A). This also allows students to work with a device that is directly relevant to a key application space for DNA origami, namely drug delivery. While it is not essential for learning the basics of DNA origami fabrication and analysis, which is the focus here, we envision the design process could be introduced in parallel as desired.

**Figure 2:**
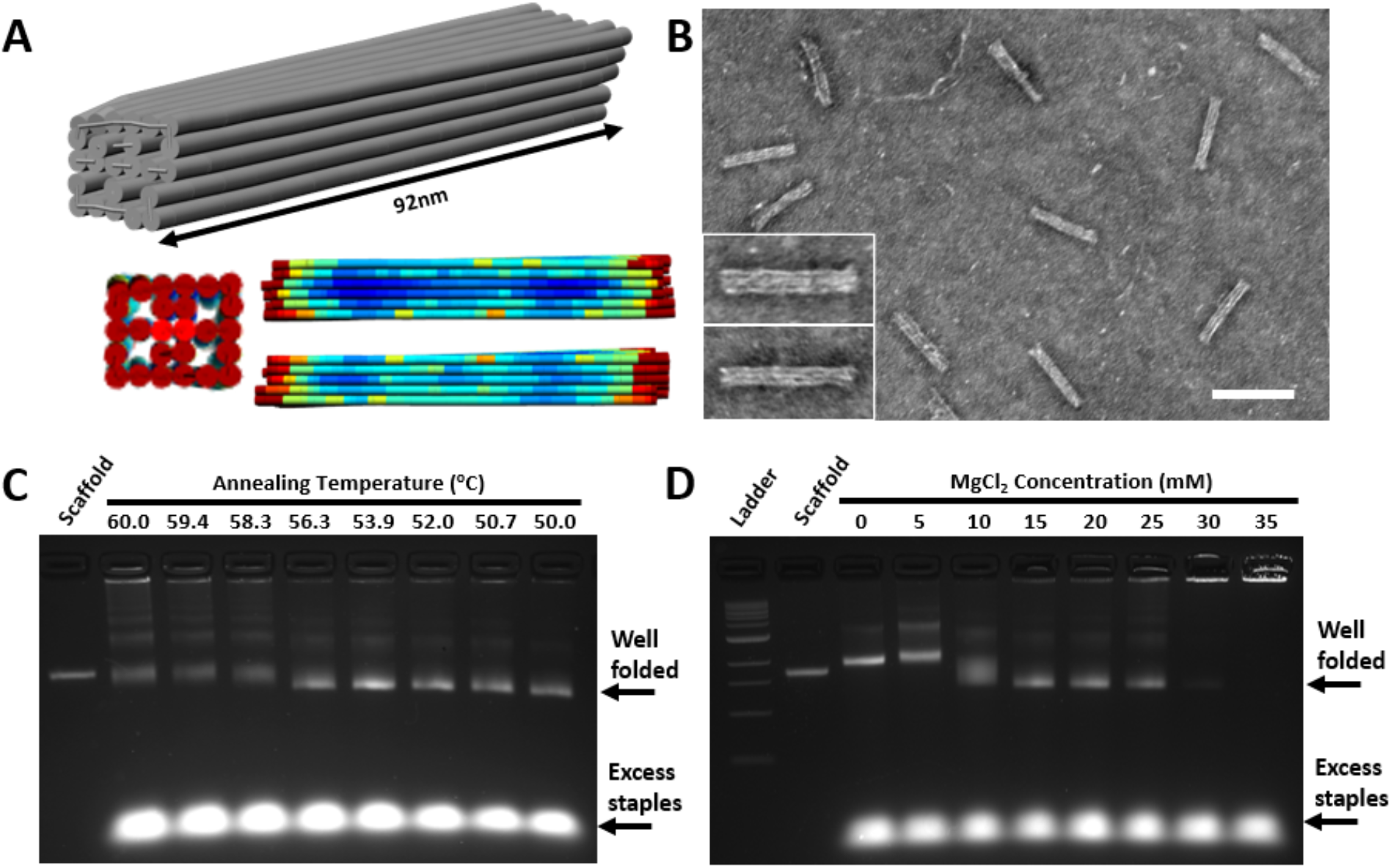
DNA origami ‘Horse’ nanorod structure design and fabrication. A) Models of the Horse nanostructure simulated using Cando^47^. The color-scale illustrates relative magnitude of root-mean-squared fluctuations. B) TEM image of the Horse (scale bar = 100 nm) folded at 20 mM MgCl_2_ in a 15 min thermal ramp with 5 min at 65°C, 15 min at 52°C, and 5 min at 4°C. Insets show zoomed in views of top and side views of the Horse nanostructure. (C) Agarose gel electrophoresis (AGE) analysis of nanostructures folded in a thermocycler for 10 minutes at varying annealing temperatures compared with the M13mp18 scaffold. The gel shows well-folded structures over a range of annealing temperatures from 50.0 - 56.3°C that run slightly faster than the scaffold and the excess staples that run much faster than the folded structures. (D) AGE analysis of nanostructures folded in a thermocycler at 52°C for 10 minutes with varying concentrations of MgCl_2_ compared with a 100 kB ladder and scaffold show well-folded structures at MgCl_2_ concentrations ranging from 15-25 mM. Structures are misfolded at 0-5 mM (indicated by slower running bands), partially folded at 10 mM (indicated by slightly slower and smeared band), and aggregate at 30-35 mM (indicated by bright signal stuck in the well).

### Rapid fabrication of DNA origami

The Horse nanostructure fabrication was originally carried out via self-assembly in a thermocycler over the course of ~17 hours. To reduce this time and eliminate the need for costly equipment, we built on recent work demonstrating faster^24,44^ and low-cost^24^ methods to fold DNA origami structures. Halley et al^24^ demonstrated the Horse structure folds well within a range of constant annealing temperatures between 40 and 60°C when annealed for four hours, and at a particular annealing temperature, the Horse structure can fold in as little as 10 min of annealing. This suggests folding the Horse structure only requires holding a melting temperature at 65°C and an annealing temperature at 52°C, followed by rapid cooling, which could be carried out using equipment such as water baths and ice buckets.

Expanding on this work, we aimed to minimize the folding time without the need for precise temperature control. We tested two critical parameters for folding, the annealing temperature and the MgCl_2_ concentration. The presence of positive ions is essential to screen the repulsions of the negatively charged phosphate groups on the DNA strands (i.e., charge screening), and temperature regulates the stability of binding interactions between the staples and scaffold allowing the strands to bind in the most stable configuration. We tested these parameters with a rapid thermal cycle consisting of 5 min at 65°C, 10 min at a constant annealing temperature (performed in a laboratory thermocycler), followed by rapid cooling to 4°C. We first tested a range of constant annealing temperatures from 60 to 50°C using a MgCl_2_ concentration of 20 mM, which leads to high yield assembly for longer annealing times^24^. The folding results were analyzed by TEM (Fig. 2B) and agarose gel electrophoresis (AGE), performed with laboratory AGE equipment (Fig. 2C). These results show that with this ~15 min folding protocol the structures fold at annealing temperatures in range of 50 to 56°C, with 56°C showing a decreased yield as indicated by the slightly smeared band. These results suggest that the Horse nanostructure folds in ~15 min with highest yields observed in the 50-54°C annealing temperature range.

We also investigated how the ~15 min folding is affected by varying MgCl_2_ concentrations, both to confirm 20 mM MgCl_2_ remains an optimal concentration and as a precursor to the intended laboratory experiment module, which focuses on introducing the concept of screening salt concentrations as a common optimization step for DNA origami fabrication. Horse nanostructures were folded in a laboratory thermocycler at 52°C with the same ~15 min thermal cycle and MgCl_2_ concentrations varied from 0 to 35 mM MgCl_2_ in 5 mM increments. AGE results (Fig. 2D) show that structures begin to form at 10 mM MgCl_2_ and fold most efficiently at 15-20 mM MgCl_2_. At 25 mM and above, structures exhibit increasing aggregation, indicated by the build-up of signal in the wells, since large aggregates cannot migrate into the gel.

For both sets of experiments we confirmed that the leading bands consisted of well-folded Horse nanostructures by TEM imaging. Fig. 1B shows a representative TEM image of Horse nanostructures folded at 20 mM MgCl_2_ using the ~15 min folding protocol with annealing at 52°C, depicting well-folded structures. Combined, these results show Horse structures can fold with high yield within ~15 min at 20 mM MgCl_2_, including 10 min of annealing at temperatures in the range of 50-54°C. While we used 65°C for melting here, prior work has used up to 95°C for the melting phase^6,7^, suggesting precise control of the melting temperature is also not critical.

### Rapid fabrication of DNA origami with classroom ready equipment

We aimed to translate this fast and simple folding approach to classroom-ready equipment. Building on Halley et al^24^, who demonstrated DNA origami folding with heated water baths, we developed a folding approach that utilizes two hot plates to heat two one-liter beakers filled with ~500 ml of water (i.e., water baths) and an ice bucket (or additional beaker) filled with ice and water. The heated water baths were used to hold a melting temperature of ~65-70°C and an annealing temperature of 52-53°C, and the ice bucket was used for the final cooling step. Temperatures were monitored using standard laboratory thermometers placed in each water bath. Rather than seeking specific hot plate settings to achieve the correct temperatures, we established a simpler approach that relies on manual control of the temperatures (Fig. 3A). Both hotplates were kept on their high (500°C) setting, and beakers were placed on the hot plate until the water baths reached the desired temperature range. The beakers were then removed from the hotplate when they reached the target temperature range and placed on the lab bench on a piece of cardboard (for some insulation). Beakers were placed back onto the hot plate when they reached the lower end of the target temperature range. With our experimental setup, we determined that placing the beaker back onto the hot plate for 2-3 seconds on the high setting would raise 500 mL of water by about 1 degree. This manual back-and-forth process between the hotplate and the cardboard was used during the duration of the 5-minute melting and 10-minute annealing steps.

**Figure 3:**
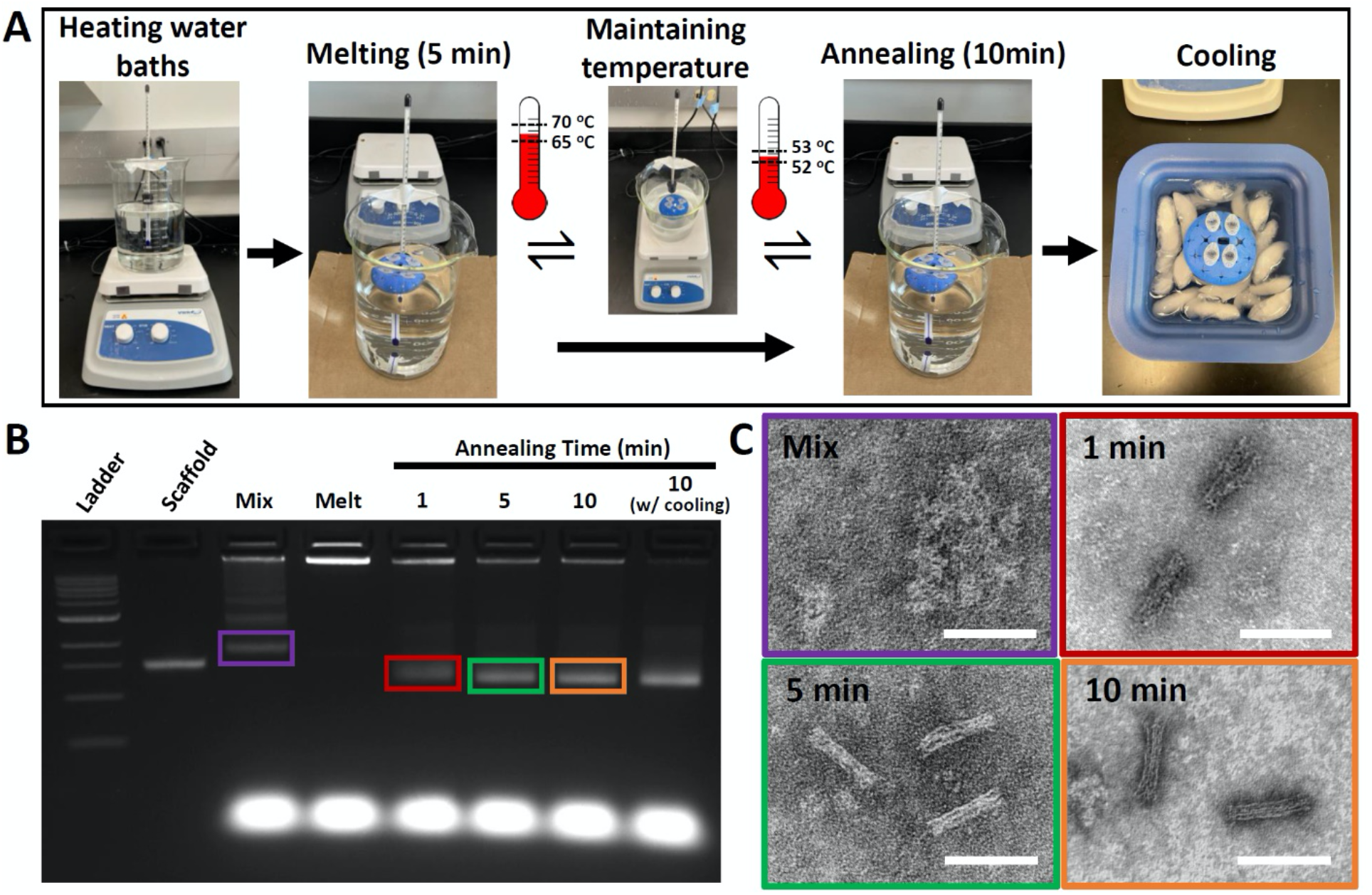
Classroom-ready fabrication of DNA origami Horse nanostructures. (A) Two water baths are prepared with 500 mL of water in a 1 L beaker. One for melting is maintained at temperatures in the range of 65-70°C and the other at the proper annealing temperature, which is 52-53°C for the Horse structure. Temperatures are monitored with thermometers directly in the heated water baths. The beakers are removed from the hot plate and placed on the lab bench on top of cardboard (for insulation) when reaching the upper limit of the temperature range and placed back on the hot plate when reaching the lower limit of the target temperature range. Folding reactions in small tubes containing DNA scaffold and staple strands are placed in the melting beaker for 5 minutes, then in the annealing beaker for 10 minutes, and finally cooled in an ice bath for 5 minutes. After the mixing stage, melting stage, and after 1,5, or 10 minutes of folding the solutions were rapidly quenched in liquid nitrogen and then thawed and subjected to B) AGE and C) TEM analysis to illustrate the progression of the fabrication process. These results illustrate structures form misfolded ‘blobs’ of DNA upon mixing. The melt phase shows aggregated DNA, which may have occurred in sample handling prior to loading the gel. Structures are already mostly folded by 1 min of annealing, and fold completely at 5 and 10 min of annealing. The final 10-minute lane on the gel was cooled in an ice bath. (Scale bars = 100 nm)

Horse nanostructure fabrication was carried out by putting the tube containing the folding reaction solution into the melt water bath for 5 minutes while it was kept in the target temperature range. Tubes were placed into water baths using foam tube holders thin enough to keep the liquid inside the tube submerged below the water level (Fig. 3A). After 5 minutes, the tubes were moved to the annealing water bath for 10 minutes while it was held in the desired temperature range. Once 10 minutes had elapsed, the tubes were transferred to an ice bath for ~5 minutes. To reveal better insight into this classroom-ready rapid folding approach, we assessed the folding reaction at several stages during the process, after mixing at room temperature, after the melting phase, and after various annealing times, by assessing a small volume of the folding reaction solution with AGE (Fig. 3B) and TEM (Fig. 3C). Melt and annealing stage samples were quenched in liquid nitrogen to ‘flash-freeze’ the reaction and loaded into gels directly after thawing. TEM images were then taken using gel purified samples. AGE revealed slower migrating DNA constructs after mixing, likely due to the scaffold binding staples without forming any well-defined structure; and DNA appeared to be aggregated at the end of the melt phase (stuck in the well), although this may have occurred during sample handling. Both AGE and TEM revealed the basic shape (although not well-defined) folds even with 1 min of annealing, and both 5 and 10 min of annealing led to well-folded structures, which is consistent with prior results^24^.

### Analysis of DNA origami folding using classroom ready equipment

Traditional laboratory electrophoresis equipment is expensive (~$1000-3000 for a gel rig including required power supply and light source for visualization), making it impractical for many K-12 or undergraduate-level science classrooms. To facilitate broader access, especially for education, some companies have developed inexpensive, safe, and portable gel electrophoresis systems. Here we implemented the MiniOne gel electrophoresis system designed for classroom use48484848, which contains a gel rig, power supply, and light source all in one compact system at significantly lower cost (<$300 kit, also includes 20 μL pipette and gel casting equipment). Figures 4A and 4B compare the equipment needed to setup and run AGE using the MiniOne and research laboratory equipment, respectively.

**Figure 4:**
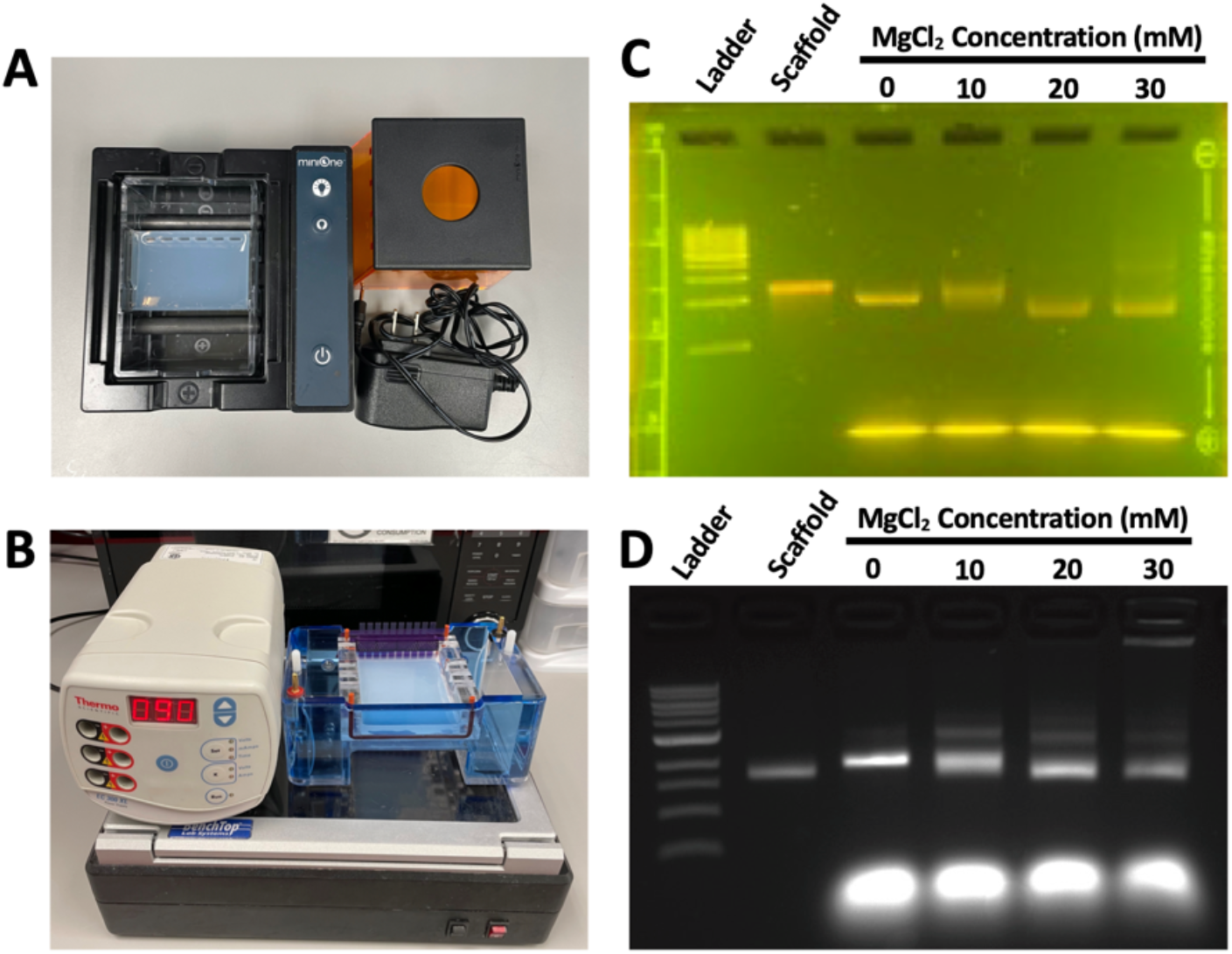
Gel electrophoresis setups. using (A) the MiniOne and (B) research laboratory equipment. AGE analysis of the folding reactions using varying levels of MgCl_2_ run alongside a 1 kB ladder and the scaffold for (C) the MiniOne and (D) research laboratory equipment. Gel shifts illustrate expected results where nanostructures are misfolded at 0-10 mM, well-folded at 20 mM, and begin to aggregate at 30 mM.

To test the MiniOne electrophoresis system for DNA origami analysis, we performed a salt screen (similar to Figure 2D) using the previously described ~15 min hot plate water bath folding approach. Since the MiniOne gels contain 6 lanes, we condensed our salt screen to include folding reactions carried out with 0, 10, 20, and 30 mM MgCl_2_, leaving room to run a DNA ladder and the ssDNA scaffold as references on the gel. This also conserves materials, which may be an important consideration for educational implementation. The results of these reactions were analyzed by AGE (Fig. 4C-D). The MiniOne, however, is designed to run AGE experiments for standard DNA analyses (i.e. not analysis of DNA origami), which are run under different than those typically used for running AGE with DNA origami. DNA origami structures are typically run on 2% agarose gels in a buffer containing 11 mM MgCl_2_, both in the gel and the surrounding running buffer because positive counterions maintain the stability of DNA origami nanostructures. However, we found the MiniOne system could not run with these conditions, likely due to the excessive current generated from the high concentration of ions, which tends to heat the gel and running buffer. This heating is normally accounted for by cooling the gels in an ice water bath while they run, but this is not practical in the MiniOne system. We tested a series of AGE conditions to converge to a protocol that was compatible with the MiniOne system, maintained DNA origami stability, and enabled visualization of gel shifts indicative of changes on DNA origami folding quality. The resulting AGE altered conditions consisted of a 1% agarose gel with 3 mM MgCl_2_ in the gel and running buffer. Importantly, the MiniOne gel kits use GelGreen stain for DNA staining, reducing the risks of exposure to the standard mutagenic stain, ethidium bromide, and harmful UV light exposure.

Figures 4C and D compare AGE analysis of the salt screen on the MiniOne setup and laboratory gel equipment, respectively. Both gels illustrate that 0-10 mM MgCl_2_ leads to misfolded structures. The Horse structure folds well at 20 and 30 mM, but 30 mM also leads to some aggregation, as indicated by the trailing smear in MiniOne gel and signal in the well on the laboratory equipment gel.

One additional advantage of the MiniOne system is that the gel migration can be viewed in realtime since the visualization is performed directly on the same system, as opposed to having to transfer the gel to an imager for the laboratory equipment. Figure 5 shows snapshots of the MiniOne salt screen gel taken at 10 min time intervals (note: 40 min is same as Fig. 4C). Structures were visualized using the built-in high energy LED light source of the MiniOne system, and images were acquired using a smart phone camera. These results illustrate the relevant gel details (gel shifts indicating well-folded structures and aggregation) can be observed after 30 minutes. We further confirmed these folding results with TEM imaging of structures purified from the MiniOne gel. Figure 5B shows representative TEM images for each MgCl_2_ concentration, confirming misfolded structures at 0 mM, partially folded structures at 10 mM, and well folded structures at 20-30 mM.

**Figure 5:**
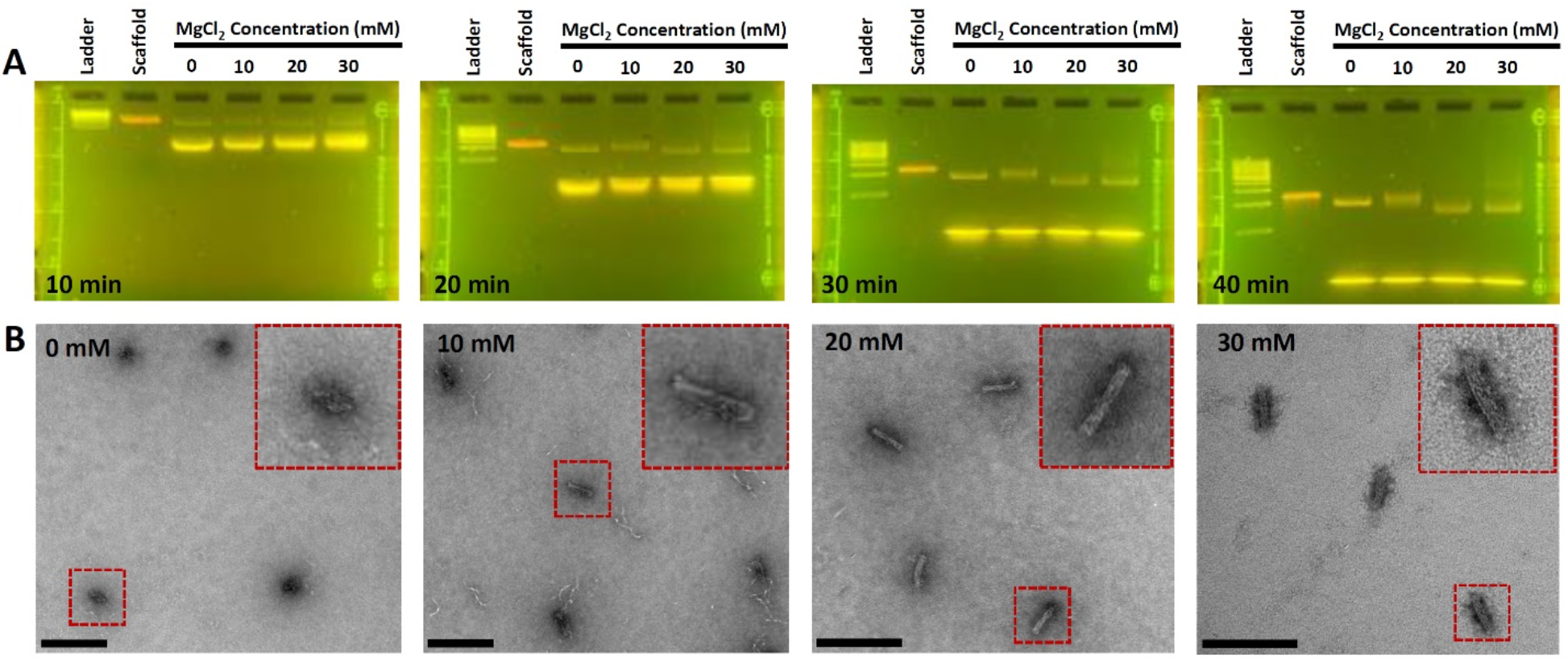
DNA origami classroom AGE analysis of MgCl_2_ salt screen. A) AGE analysis of Horse nanostructures folded at several MgCl_2_ concentrations was carried out on the MiniOne system and imaged at several timepoints. The relevant gel shift indicating proper folding at 20 mM and aggregation at 30 mM are noticeable by 30 min and clearly visible by 40 min. This MiniOne AGE salt screen represents a folding analysis that can be carried out in a classroom. B) TEM images of sample purified from the MiniOne gels confirm poor folding at 0 mM, partial folding at 10 mM, and high-quality folding at 20-30 mM. Scale bars = 200 nm.

## ADAPTATION TO THE CLASSROOM

Time, cost, and complexity are three important factors when developing the protocol to perform this experiment in middle school, high school, and undergraduate classrooms. We have shortened the time required to perform the experiments by developing protocols that can be completed in ~2 hr, or two one-hour sessions. Alternatively, these experiments could be performed in a shorter ~one-hour single session with additional instructor setup and preparation (casting of gels, bringing water baths up to temperature, pre-mixing components of folding reactions, etc.). We also reduced costs by eliminating the need for specialized equipment including the thermocycler (~$5,000-$10,000) and laboratory gel electrophoresis equipment (~$1000-2000), UV table and imager (~$5000-$10,000) and replacing these with simpler, cheaper, and possibly readily available equipment consisting of hot plates (~$200-300 if not available), glassware (~$20 per beaker, 2 needed), and the MiniOne gel electrophoresis system (~$300). A detailed cost comparison with specific examples is provided in Supplemental Table 1. We also significantly reduced complexity by streamlining the folding and analysis process and using the compact classroom-ready equipment.

**Table 1.** Required equipment and reagents.

| Equipment |  | Reagents |  |
| --- | --- | --- | --- |
| E1 | Scale | R1 | Distilled water |
| E2 | Chemical Spoon | R2 | MgCl<sub>2</sub> |
| E3 | Beakers | R3 | GelGreen DNA stain |
| E4 | Microwave | R4 | Agarose |
| E5 | Timer | R5 | 0.5x TBE |
| E6 | Floating tube rack | R6 | 1 kb DNA ladder |
| E7 | Graduated cylinder | R7 | Loading Dye |
| E8 | Hotplate | R8 | Folding Reaction Buffer |
| E9 | Thermometer | R9 | M13mp18 DNA scaffold |
| E10 | Gloves | R10 | Horse oligos |
| E11 | Calulator |  |  |
| E12 | Eppendorf tubes |  |  |
| E13 | Pipette and Tips |  |  |
| E14 | Gel Casting Equipment |  |  |
| E15 | Electrophoresis System |  |  |

Here we lay out a proposed procedure for a ~2 hr experiment module for classroom implementation of the DNA origami salt screen presented in Figure 4A. The procedure consisting of three main steps: 1) Preparing the Gel for Electrophoresis, 2) Running the Folding Reaction, and 3) Running the Gel. Each step entails preparation time (which will vary based on the students’ prior experience with lab work (pipetting, measuring reagents, etc.)), and each step has a rate limiting step described below.

### Step 1 - Preparing the Gel for Electrophoresis (~35 min)

Students will prepare the gel running buffer and cast the gel. This involves both standard laboratory measurements as well as pipetting. Preparing the gel takes approximately fifteen minutes. The rate limiting step here is waiting for the gel to solidify, which takes ~30 min at room temperature. Students can prepare the folding reaction or bring water baths up to target temperatures while waiting for the gel to solidify.

### Step 2 - Running the Folding Reaction (~30 min)

In this step, students will prepare the two water baths bringing them to the desired target temperature range, mix the folding reaction, and perform the folding thermal cycle. Students will need to calculate appropriate dilutions to make the different 10x concentrations of a salt buffer (0, 100, 200, and 300 mM MgCl_2_), and then mix the five ingredients at proper volumes and concentrations (scaffold, staple strands, folding buffer, salt buffer, and water). This process takes ~ 15-20 min. The rate limiting step is folding the structure for 20 minutes (5-min melt, 10-min fold, and 5-min cooling).

### Step 3 - Running the Gel (~50 min)

Here, students will perform gel electrophoresis by setting up the gel equipment, mixing the folded structure solution with gel loading dye, loading the samples into wells, and running the gels. Preparation takes ~10 min, and running the gel takes 30-40 min to visualize the gel shift results. Students will compare their results to expected results shown in Figures 4C or 4D, depending on the electrophoresis setup being used.

The total protocol can be completed in a single, two-hour lab session. If broken up into two one-hour sessions, the initial session would consist of Step 1 and the preparation for Steps 2 (i.e., mixing folding reactions). Session two would then include running the folding reaction from Step 2 as well as running the gel in Step 3. Part of the preparation of Step 3 (i.e., preparing loading dye) can be done during the folding reaction, as long as students can also carefully monitor the water bath temperatures simultaneously. The protocol can also be completed in a single one-hour session if the instructor prepares the gel, water baths, and folding reaction mixtures ahead of time. Students would then perform the folding reaction, mix with loading dye, load and then run the gel for ~30 min. This shorter method would be ideal for younger students or students with no prior lab experience.

Table 1 shows the equipment and reagents needed to complete the entire procedure. Many items on the equipment/supplies list (E1-E12) are commonly found in classroom science laboratories or can be purchased at a low cost. E13-E15 are not as common, however, inexpensive classroom versions exist such as the MiniOne Gel Electrophoresis Kit used in this research (< $300).

The reagents R1 and R2 are readily found in science laboratories. R3-R7 can also be purchased at low cost individually or in kits (in these experiments, reagents R3-R7 were purchased from MiniOne). The reagents R8-R10 can be provided in small quantities for those interested in performing the procedure.

## CONCLUSION

In this work, we developed a classroom-ready approach to folding DNA origami nanostructures, and we demonstrated a protocol to perform an experiment that analyzes the effect of salt concentration on DNA origami folding, which is a common first step in optimizing fabrication. These procedures can be done in a time and cost-effective manner, and variation in complexity allows this to be a valuable educational experience for middle school, high school, and undergraduate science students in a variety of disciplines. To date, either the extended or condensed version of these experiments have been completed with middle and high school teachers at the Science Education Council of Ohio (SECO) conference 2019 and at the Association for Biology Laboratory Education (ABLE) workshop 2018. The condensed (~1hr) experiment was also performed with middle school students at Hilltonia middle school. The full experiment has also been carried out in undergraduate classrooms both at OSU in a Mechanical Engineering course (~20 students) and by two different classes of Systems and Mechanical Engineering students (~50 students) at Otterbein University.

This work is a foundation for implementing and translating DNA nanotechnology education with a hands-on approach to classrooms for undergraduate, secondary, and primary school students. Previous efforts translating DNA nanotechnology to undergraduate classrooms demonstrated DNA nanoswitches to introduce concepts of biosensing applications and conformational changes of DNA constructs^49^. This work expands upon this foundation by introducing the highly versatile and widely applicable DNA origami nanotechnology. We envision this can stimulate additional opportunities to translate additional concepts and functions of DNA origami to classrooms such as dynamic or complex DNA origami nanostructures and design and simulation modules. Translating DNA origami nanotechnology into classrooms will play an important role in exposing young students to this highly promising cutting-edge approach that is likely to impact a wide range of industries, which can educate students about potential STEM-related fields and careers and reinforce other fundamental science and engineering learning milestones.

## Supporting information

List of DNA nanostructure staples

## ACKNOWLEDGMENTS

This work was supported by start-up funding provided to MWH by Otterbein University and by NSF grants #1916740, #1921881, and #1351159 to CEC. We thank the Otterbein University Department of Engineering for purchasing the electrophoresis equipment used in the initial classroom implementations and to optimize protocols. We would also like to thank the Otterbein Engineering classes of 2019 and 2021 for testing the protocols and providing valuable feedback and members of the Castro Laboratory for providing useful feedback during development of the protocols. We would also like to acknowledge the OSU Pelotonia fellowship program for funding PEB. Transmission electron microscopy images were acquired at the OSU Campus Microscopy and Imaging Facility, which is supported in part by grant number P30 CA016058, National Cancer Institute, Bethesda, MD.

## ASSOCIATED CONTENT

### Supporting Information

Supplemental material (PDF)

Horse Staple Sheet (XLSX)

## Footnotes

1 The British chemist, Rosalind Franklin, also played a central role in understanding the structure of DNA and viruses, laying the foundation for structural virology^27,28^.

