## Supplemental tables and figures for "Translating DNA origami Nanotechnology to Middle School, High School, and Undergraduate Laboratories"

**Supplemental Material**

Table S1: A comparison between the costs of the fabrication and the evaluation of DNA origami structures in a research laboratory vs our proposed classroom demo. Research lab estimates are conservative, not taking in consideration readily available supplies, such as PCR tubes and other plasticware. For comparison, we selected lab equipment at the low end of the cost range for the classroom demo.

| **Process** | **Research Lab Setting** | | **Classroom Demo** | |
| --- | --- | --- | --- | --- |
|  | **Equipment** | **Cost** | **Equipment** | **Cost** |
| **Folding** | Thermocycler (BioRad-T100) | $5367 | Hotplate (Thermo Scientific Cimarec Basic) | $229 |
|  | P20 pipette (Fisherbrand) | $337 | P20 pipette (Included with MiniOne Electrophoresis System) | $0 |
| **Gel Electrophoresis** | Microwave | $100 | Microwave | $100 |
|  | Fisherbrand Minigel Horizontal Electrophoresis System | $398 | MiniOne Electrophoresis System | $280 |
|  | Power Supply (Fisherbrand-FB300) | $890 |  |  |
|  | Imaging (UVP PhotoDoc-It - UVP  97027404) | $5300 |  |  |
| **Total Cost** | **$12392** | | **$609** | |

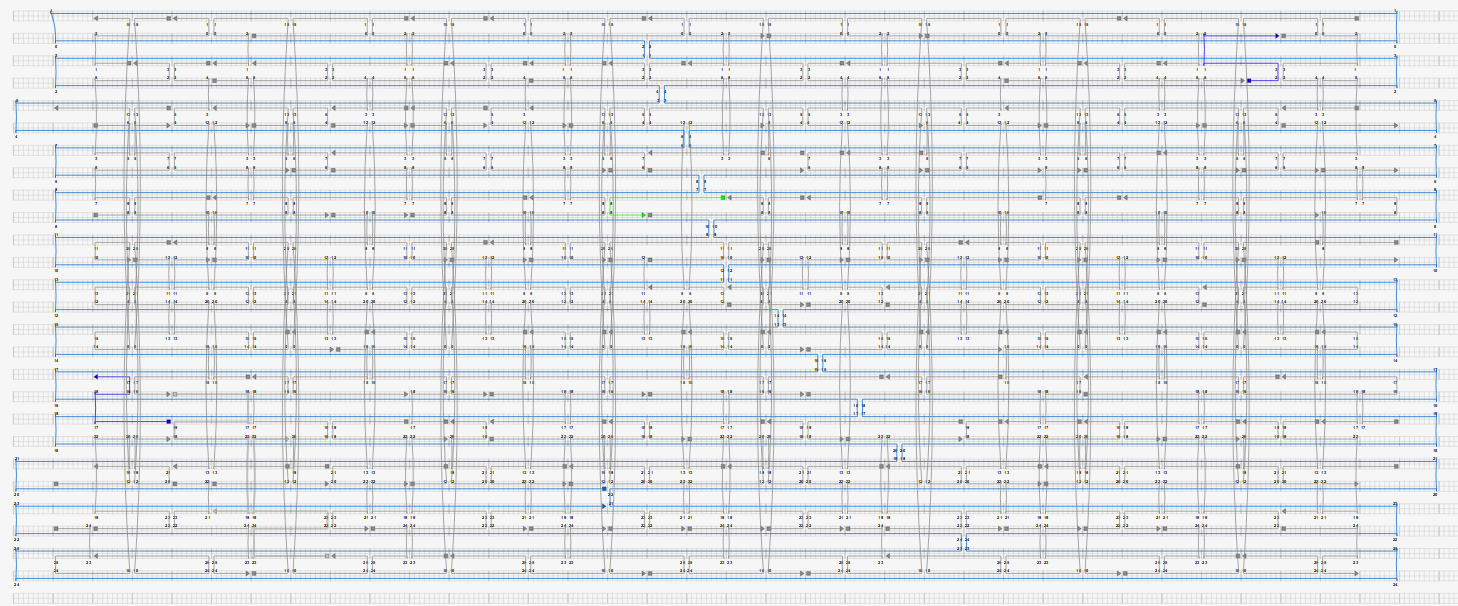

Figure S1: Cadnano design of the Horse structure.

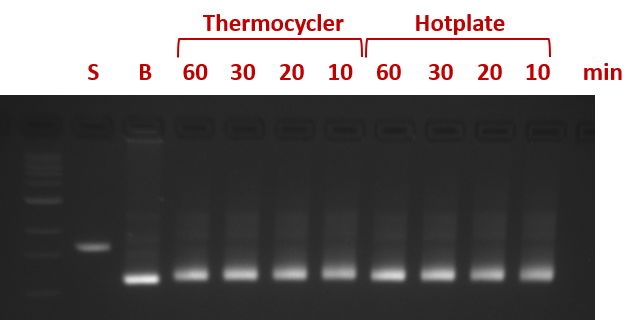

Figure S2: Proof-of-concept gel to show that the structure can be folded at a single temperature (at 20 mM salt) as a function of time on hotplate by directly comparing to identical conditions folded on a thermocycler.

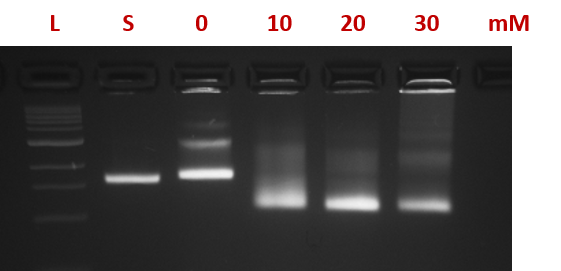

Figure S3: Gel of varying salt concentrations using a 2%, 11 mM MgCl_2_ gel.
